# Sentinel Cards Provide Practical SARS-CoV-2 Monitoring in School Settings

**DOI:** 10.1101/2022.02.01.478759

**Authors:** Victor J Cantú, Karenina Sanders, Pedro Belda-Ferre, Rodolfo A Salido, Rebecca Tsai, Brett Austin, William Jordan, Menka Asudani, Amanda Walster, Celestine G. Magallanes, Holly Valentine, Araz Manjoonian, Carrissa Wijaya, Vinton Omaleki, Stefan Aigner, Nathan A Baer, Maryann Betty, Anelizze Castro-Martínez, Willi Cheung, Peter De Hoff, Emily Eisner, Abbas Hakim, Alma L Lastrella, Elijah S Lawrence, Toan T Ngo, Tyler Ostrander, Ashley Plascencia, Shashank Sathe, Elizabeth W Smoot, Aaron F Carlin, Gene W Yeo, Louise C Laurent, Anna Liza Manlutac, Rebecca Fielding-Miller, Rob Knight

## Abstract

Accurate, high-resolution environmental monitoring of SARS-CoV-2 traces indoors through sentinel cards is a promising approach to help students safely return to in-person learning. Because SARS-CoV-2 RNA can persist for up to a week on several indoor surface types, there is a need for increased temporal resolution to determine whether consecutive surface positives arise from new infection events or continue to report past events. Cleaning sentinel cards after sampling would provide the needed resolution, but might interfere with assay performance. We tested the effect of three cleaning solutions (BZK wipes, wet wipes, RNase Away) at three different viral loads: “high” (4 x 10^4^ GE/mL), “medium” (1 x 10^4^ GE/mL), and “low” (2.5 x 10^3^ GE/mL). RNAse Away, chosen as a positive control, was the most effective cleaning solution on all three viral loads. Wet wipes were found to be more effective than BZK wipes in the medium viral load condition. The low viral load condition was easily reset with all three cleaning solutions. These findings will enable temporal SARS-CoV-2 monitoring in indoor environments where transmission risk of the virus is high and the need to avoid individual-level sampling for privacy or compliance reasons exists.

**Importance:** Because SARS-CoV-2, the virus that causes COVID-19, persists on surfaces, testing swabs taken from surfaces is useful as a monitoring tool. This approach is especially valuable in school settings, where there are cost and privacy concerns that are eliminated by taking a single sample from a classroom. However, the virus persists for days to weeks on surface samples, so it is impossible to tell whether positive detection events on consecutive days are persistent signal or new infectious cases, and therefore whether the positive individuals have been successfully removed from the classroom. We compare several methods for cleaning “sentinel cards” to show that this approach can be used to identify new SARS-CoV-2 signals day to day. The results are important for determining how to monitor classrooms and other indoor environments for SARS-CoV-2 virus.

## Body

For the last two years, the SARS-CoV-2 pandemic has disrupted lives and caused millions of deaths globally. Due to the high risk of virus transmission in indoor settings, schools have been forced to convert to remote learning [1]. Although remote learning can be convenient for some, not every child has access to a stable internet connection and a supportive, quiet learning environment [2,3]. Therefore, most child health authorities are recommending a return to in-person learning, if it can be conducted safely [4]. Effective SARS-CoV-2 monitoring is crucial to allow for in-person learning to resume safely and widely [5], with the goal of restoring education equity. However, performing daily nasal swabs to monitor the spread of the disease has high financial and labor costs, and often runs into difficulties with consent and reporting of results to relevant public health authorities.

Wastewater and environmental monitoring strategies have been developed [6–8] and implemented [9] as a means of circumventing clinical swabs. We have already demonstrated that viral signals from COVID-19 patients in indoor environments commonly accumulate on high-touch surfaces and the floors in front of features with high interaction times [8]. Additionally, SARS-CoV-2 RNA has been demonstrated to persist for up to a week on several indoor surface types [7, 10], making it difficult to understand exactly when an infected individual came into contact with a surface or if consecutive positives are from new deposition events. Thus, an effective post-sampling cleaning procedure needs to be established in order to increase temporal resolution and ensure that consecutive positives are from new infection events.

To increase the temporal resolution of proven environmental pipelines [9,11] we tested resetting SARS-CoV-2 RNA signal with a mock sentinel surface. Here, a sentinel surface is a surface used as an environmental monitoring tool for detecting whether or not an infected individual was recently present in an indoor space. The mock sentinel surfaces we used were 100 cm^2^ laminated cards. The sentinel cards were inoculated with 10 μL of a dilution series of heat-inactivated SARS-CoV-2 particles (strain WA-1, SA-WA1/2020) in water and then wiped with a cleaning solution each day for five days. Samples were collected by swabbing the sentinel cards pre-inoculation, post-inoculation, and post-wipe (Supplemental Fig. S1).

For this study we used three viral loads: “high” (4 x 10^4^), “medium” (1 x 10^4^), and “low” (2.5 x 10^3^) dilutions of SARS-CoV-2 viral genomic equivalents, as measured by droplet digital PCR. These concentrations were chosen to bracket the ranges we typically observed in classrooms during SASEA [9]. We used two different transport media: SDS (0.5% w/v sodium dodecyl sulfate (SDS), Acros Organics, 230420025), which we have previously shown to yield superior results in SARS-CoV-2 molecular assays [County paper], and VTM (Viral Transport Medium, NEST Scientific USA, 202016), which is in widespread use by public health laboratories. We tested three cleaning methods: benzalkonium chloride (BZK) antiseptic towelettes (Dynarex, 1331), moist wet wipe (WW) towelettes (Royal, RF1MB), and paper towels moistened with RNase AWAY (RA) (ThermoFisher Scientific, 10328011).

To continue benchmarking proven environmental pipelines [7, 9, 11] and to account for potential interactions, we used a factorial study design covering two swabbing media (SDS, VTM), three cleaning solutions (BZK wipes, wet wipes, RNase Away) and three viral spike-in concentrations (High, Medium, Low). Each condition was performed in triplicate for a total of 54 cards. A three-step swabbing process was performed on each card over a five-day period. First, we swabbed each card at the start of the day (Step 1). Next, the viral spike-in was added to the card and a second swab was collected (Step 2). The card was then wiped with the cleaning solution and a final swab was collected (Step 3). Extraction and RT-qPCR were performed as described in our previous work, with VTM samples processed by the Perkin-Elmer pipeline and SDS samples processed by the Thermo pipeline described in that work [PHL paper].

Our results demonstrated that all of the cleaning methods worked well at low viral load over 5 cleaning cycles, although cleaning failures were somewhat more frequent with BZK (Fig. 1). Wet wipes and BZK performed well with SDS at medium viral loads, but only wet wipes performed well with VTM under these conditions. At high viral loads, only the combination of RNase away and SDS was able to remove the signal. Therefore, we recommend that if high viral loads (Cq < 30, with SDS) are detected on a sentinel card, that the sentinel card be replaced at the next opportunity rather than cleaned. Repeat cleaning did not degrade the sentinel card surface or the ability to detect signal. As expected from our past work [11], SDS returned lower Cq values (better signal) than VTM on the same samples.

**Figure 1.**
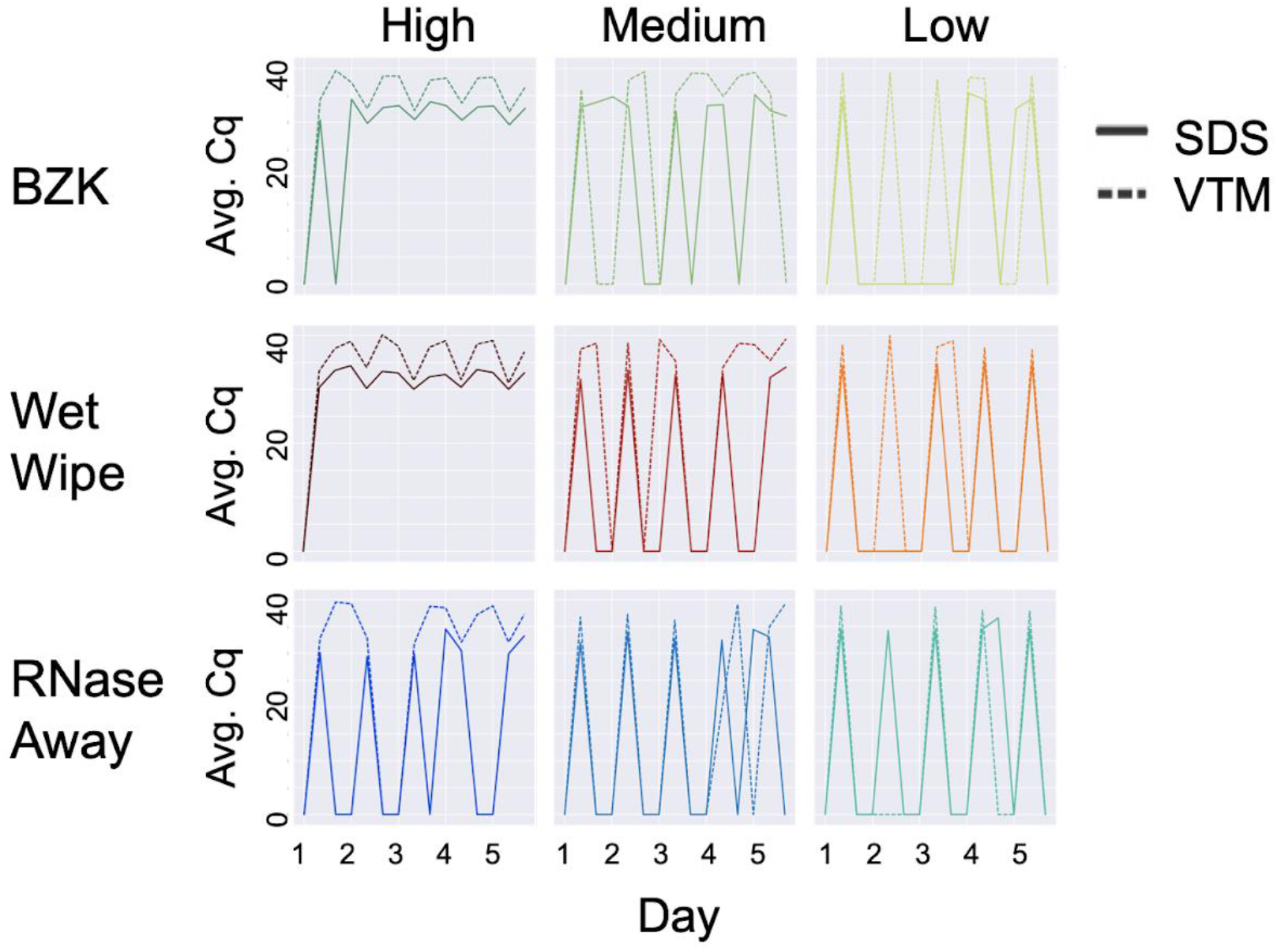
Effect of cleaning solution at high, medium and low viral load with different swabbing media. On each day, three samples were taken: (1) before addition of viral particles, (2) after addition, and (3) after cleaning. Therefore, the expected pattern is a train of 5 spikes, starting at zero, rising to the maximum Cq value, returning to zero the same day, and staying at zero until the next day, as seen for SDS in the low load condition with RNase away (bottom right panel, solid lines). High, medium, and low viral load were defined as (4 x 10^4^), (1 x 10^4^), and (2.5 x 10^3^), respectively. Average Cq (Avg. Cq) was calculated as a mean Cq value from three samples. Two viral transport media were tested: SDS (0.5% w/v sodium dodecyl sulfate (SDS) and VTM (Viral Transport Medium). Effective cleaning reset Cq for each day. RNase away was shown to be effective at each viral load, whereas benzalkonium chloride (BZK) and wet wipes were only effective at medium and low viral load.

An important consideration is the number of distinct genes recovered as matching in the RT-qPCR process, as this can make the difference between a sample being called as SARS-CoV-2 positive versus invalid. Because the peaks with the same viral load applied were highly reproducible across multiple days (reaching the same height in Fig. 1), for this analysis we could treat each day as a replicate of the pre-application, post-application, and post-cleaning sample conditions that were collected on each day. Fig. 2 shows the reproducibility of replicates with cleaning, including the number of genes amplified. Under low load conditions, as expected, cleaning was effective and non-zero values occurred nearly always post-application and disappeared on cleaning, with the exception of VTM samples which sometimes carried over (right hand column in Fig 2). In contrast, in the high load condition (left hand column in Fig. 2), cleaning was nearly always ineffective except with RNase Away, not practical for classroom use. In the medium condition (middle column), all cleaning methods were effective with SDS but none were effective with VTM – the slightly higher cluster of Cq values are obtained with VTM in each case, consistent with expectations and with Fig. 1. Because VTM is viscous and contains fetal calf serum, a noticeable film developed on the sentinel cards, and we suspect that vigorous and repeated cleaning beyond what is achievable with wipes may be required to remove all of it.

**Figure 2.**
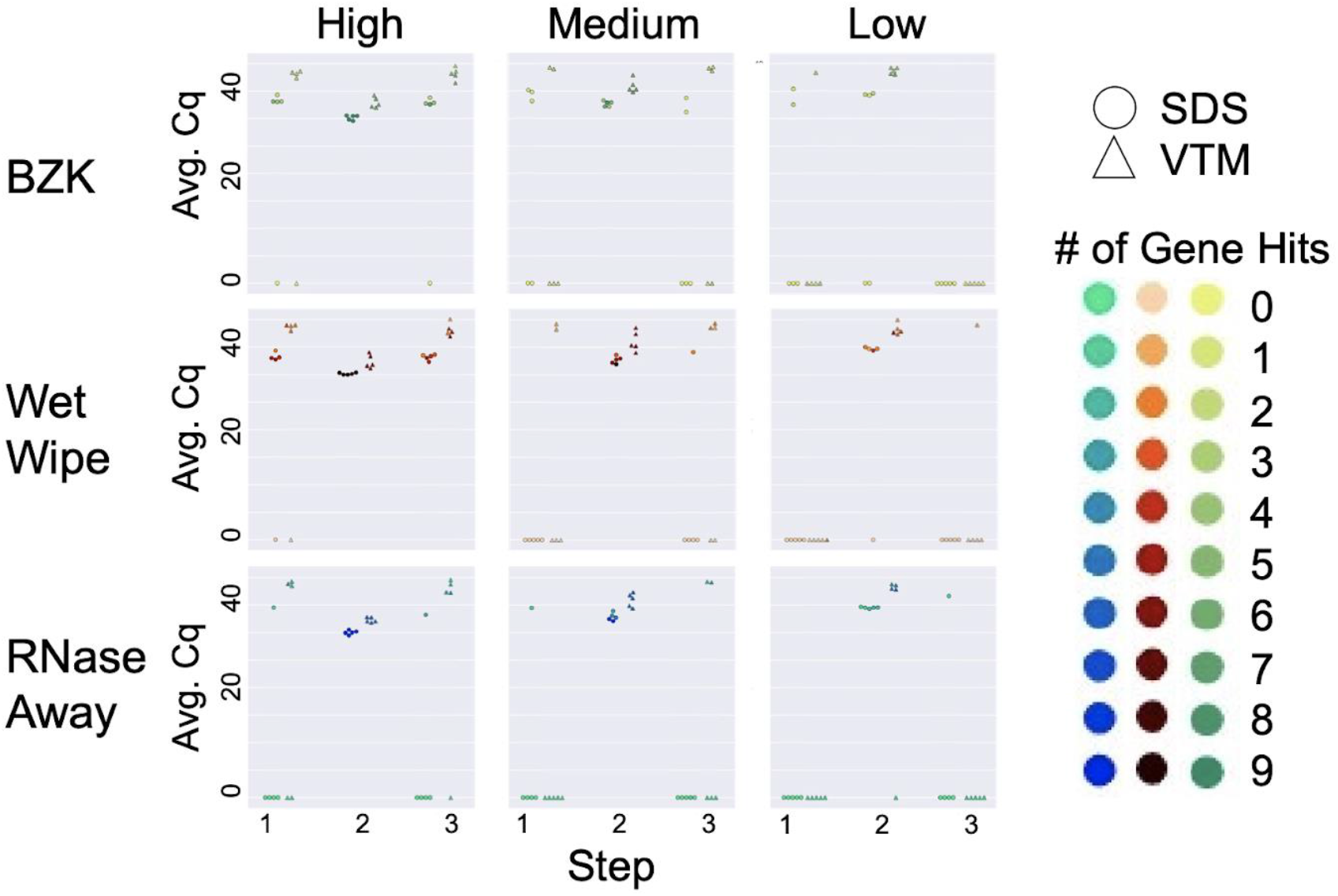
Cleaning solution efficiency after deliberate addition of viral load. Sampling was performed in three steps: initial virus amount (blank) was sampled from the wall for Step 1. Virus was deliberately loaded on the surface and sampled for Step 2. The surface was cleaned with different cleaning methods and sampled for qPCR analysis for Step 3. High, medium, and low viral load were defined as (4 x 10^4^), (1 x 10^4^), and (2.5 x 10^3^), respectively. Average Cq (Avg. Cq) was calculated as a mean Cq value from three samples. Two viral transport media were tested: SDS (0.5% w/v sodium dodecyl sulfate (SDS) and VTM (Viral Transport Medium). Effective cleaning reset Cq for each day (steps 1 and 3), whereas ineffective cleaning retained high viral load (non-zero Cq) at these steps. The number of gene hits refers to how many gene targets were amplified during RT-qPCR across the triplicate samples: the qPCR method for the SDS samples targeted 3 genes for a total of 9 possible genes amplified while the method for the VTM samples targeted 2 genes for a total of 6 possible gene hits.

Taken together, these results indicate that sentinel cards are an effective and practical solution for SARS-CoV-2 classroom monitoring, but that they must be cleaned carefully in order to remove carryover signal, and this process is easier with samples collected in SDS than in VTM (although cleaning with VTM is still possible). Because removing high viral load from sentinel cards is challenging, strong positives should be removed rather than cleaned. These findings are an important step to deployment of these cards at scale in projects such as SASEA.

## Acknowledgements

This research was supported by NIH grant (K01MH112436) to RFM, the County of San Diego Health and Human Services Agency (Contract 563236), and the Career Award for Medical Scientists from the Burroughs Wellcome Fund to A.F.C. We thank Marisol Chacon, Evelyn S Crescini, Bhavika Kapadia, Sydney C. Morgan, Alhakam Nouri, Christopher A. Ruiz, Phoebe Seaver, and Lizbeth Franco Vargas for their support with environmental SARS-CoV-2 detection as part of the EXCITE Lab.

The following reagent was deposited by the Centers for Disease Control and Prevention and obtained through BEI Resources, NIAID, NIH: SARS-Related Coronavirus 2, Isolate USA-WA1/2020, NR-52281.

---

Supplemental Fig. S1

Diagram of sampling events for each day of the experiment. Each day the sentinel cards were swabbed pre- and post-inoculation and post wiping.

